# Aquatic microbial community is partially functionally redundant: insights from an *in situ* reciprocal transplant experiment

**DOI:** 10.1101/2020.02.01.930198

**Authors:** Kshitij Tandon, Min-Tao Wan, Chia-Chin Yang, Shan-Hua Yang, Bayanmunkh Baatar, Chih-Yu Chiu, Jeng-Wei Tsai, Wen-Cheng Liu, Chen Siang Ng, Sen-Lin Tang

## Abstract

Microbial communities are considered to be functionally redundant, but few studies have tested this hypothesis empirically. In this study, we performed an *in situ* reciprocal transplant experiment on the surface and bottom waters of two lakes with disparate trophic states and tracked changes in their microbial community and functional attributes for 6 weeks using high-throughput sequencing and functional approaches. The communities from both lakes were resistant to changes in composition after the reciprocal transplant, but their functions tended to become similar to the incubating lakes’ functional profiles. A significant linear positive relationship was observed between the microbial community and functional attributes, though with varying scales of similarity, suggesting partial functional redundancy. Furthermore, the entropy-based *L*-divergence measure quantified the scale of partial functional redundancy in the lakes’ surface and bottom waters. This study establishes and quantifies the scale of partial functional redundancy in the freshwater ecosystem through empirical investigation.

## Introduction

In recent years, attempts have been made to understand the community functional relationships across different ecosystems, ranging from oceans^1–3^ to freshwater ecosystems^4,5^, soil^6,7^, plants^8,9^, and the human gut^10,11^. These studies have put forward two plausible scenarios delineating community functional relationships: 1) community and function are coupled so that changes in the community composition will affect microbe-mediated processes, and 2) where community and function appear decoupled, microbe-mediated processes are somewhat independent of changes in the community composition^12^. The latter is also known as functional redundancy.

Recently, the concept of functional redundancy has been categorized into strict and partial functional redundancy. Strict functional redundancy implies that different taxa or phylotypes can conduct the same set of metabolic processes and readily replace one another^12–14^, while partial functional redundancy implies that certain organisms can co-exist while sharing specific functions, but differ in other ecological functions or requirements^15^. The paradigm of strict functional redundancy does not hold in all ecosystems, as recently shown in soil^16,17^ and oceans^3^.

Studies of aquatic ecosystems, especially rivers and lakes, *in situ*^18–20^ and under laboratory setttings^21–23^ have been central for deducing community functional relationships; some of these studies support the presence of functional redundancy^5,24–27^, while others challenge it^28–30^. Langenheder and colleagues^25^ used laboratory bioreactors to conclude that communities with different microbial compositions may have similar functions. Delgado-Baquerizo et al.^30^ used a microcosm experiment and concluded that a decline in microbial diversity has direct effects on microbe-mediated processes. In contrast to these *in vitro* experiments, Comte et al.^31^ performed *in situ* reciprocal transplants and linked functional redundancy with metabolic plasticity in an aquatic environment using carbon metabolism as a proxy. It is important to note that differing conclusions on functional redundancy can be the results of several factors, including the sensitivity of molecular approaches, range of functions analyzed, and phylogenetic traits depths of microbes^32^. While these studies are consequential, they are limited to a few days of experimentation and focus on very broad functions (respiration, biomass replication, specific growth rate, etc.). Therefore, these and various other studies^33,34^ are limited in scope, and thus cannot provide a holistic perspective on the community functional relationships in the natural environment. Consequently, to test the functional redundancy hypothesis, a long-term *in situ* empirical study is required using a combination of high-throughput molecular and functional approaches.

In this study, we performed an *in situ* reciprocal transplantation experiment for six weeks by exchanging and incubating water from the surface and bottom of two lakes with disparate trophic states via dialysis tubes (which acted as microbial cages)^4^. We hypothesized that our pulse disturbance due to cross-transplantation and incubation would result in any of the following outcomes: 1) the microbial communities and their consequent functional profiles are both altered; 2) neither the communities nor functional profiles are altered; 3) the microbial communities are altered, but there is no change in the functional profiles; and 4) the functional profiles are altered, but there is no change in the microbial communities. These outcomes would result in the following conclusions: 1) there is no functional redundancy in the microbial communities, 2) the disturbance is not significant enough to cause any change, 3) there is functional redundancy in the microbial communities, and 4) the microbial communities are more resistant/resilient to change than are the functional profiles. A combination of meta-barcoding approaches via the 16S rRNA gene and 16S rRNA gene transcripts along with whole-metagenome high-throughput sequencing and community level physiological profiles were used to test these outcomes.

## Materials and Methods

### Sampling sites

Two proximate (21 km), sub-alpine lakes in northern Taiwan were used as experimental sites: Tsuei-Feng Lake (T) (24° 30’ 52.5’’, 121° 36’ 24.8’’ E) and Yuan-Yang Lake (Y) (24° 34’ 33.6’’ N, 121° 24’ 7.2’’ E), located at an altitude of 1,840 and 1,670 m, respectively. These lakes, despite being at similar altitudes and geographical locations, have different trophic states—Tsuei-Feng is oligotrophic^35^ and Yuan-Yang is mesotrophic^36^. Both lakes are within a protected area and are therefore exposed to minimal or no anthropogenic activity year-round. Furthermore, these lakes have been at the focus of limnological and meteorological research in Taiwan over the years.

### Experimental layout and sample collection

We conducted a reciprocal transplant experiment by which water samples from the surface (0.5 m deep) and bottom (1 m above the sediment) of each lake were transplanted and incubated for 8 weeks. The samples were collected every two weeks after transplantation (January–February 2015; three time points total). Samples were collected at the center of the lake using a boat. 2 L of additional lake water (BG) (Fig. 1A) was collected at each time point, including one at the time of transplant setup and used as controls to represent undisturbed communities and their functional profiles during the experiment. To create an artificial disturbance (for both the surface and bottom), 6 L water each from the surface and bottom of both lakes was collected, swapped, and incubated using dialysis tubes (Sigma-Aldrich, St. Louis, MO, USA) in either the same (self-swap) or a different lake (cross-swap). Specifically, T lake’s surface water was swapped and incubated in T (self-swap: Ts(n)→T) and Y lakes’ surface waters (cross-swap: Ts(n)→Y), T lake’s bottom water was swapped and incubated in T (self-swap: Tb(n)→T) and Y lakes’ bottom waters (cross-swap: Tb(n)→Y) (Fig. 1B and 1C). Self-swap samples acted as controls to monitor the effect of dialysis tubing. These dialysis tubes allow only ions (molecular weight <14 KDa) to pass through; each tube carried ~500 mL water and was placed in specially designed metal cages throughout the experiment.

**Fig 1.**
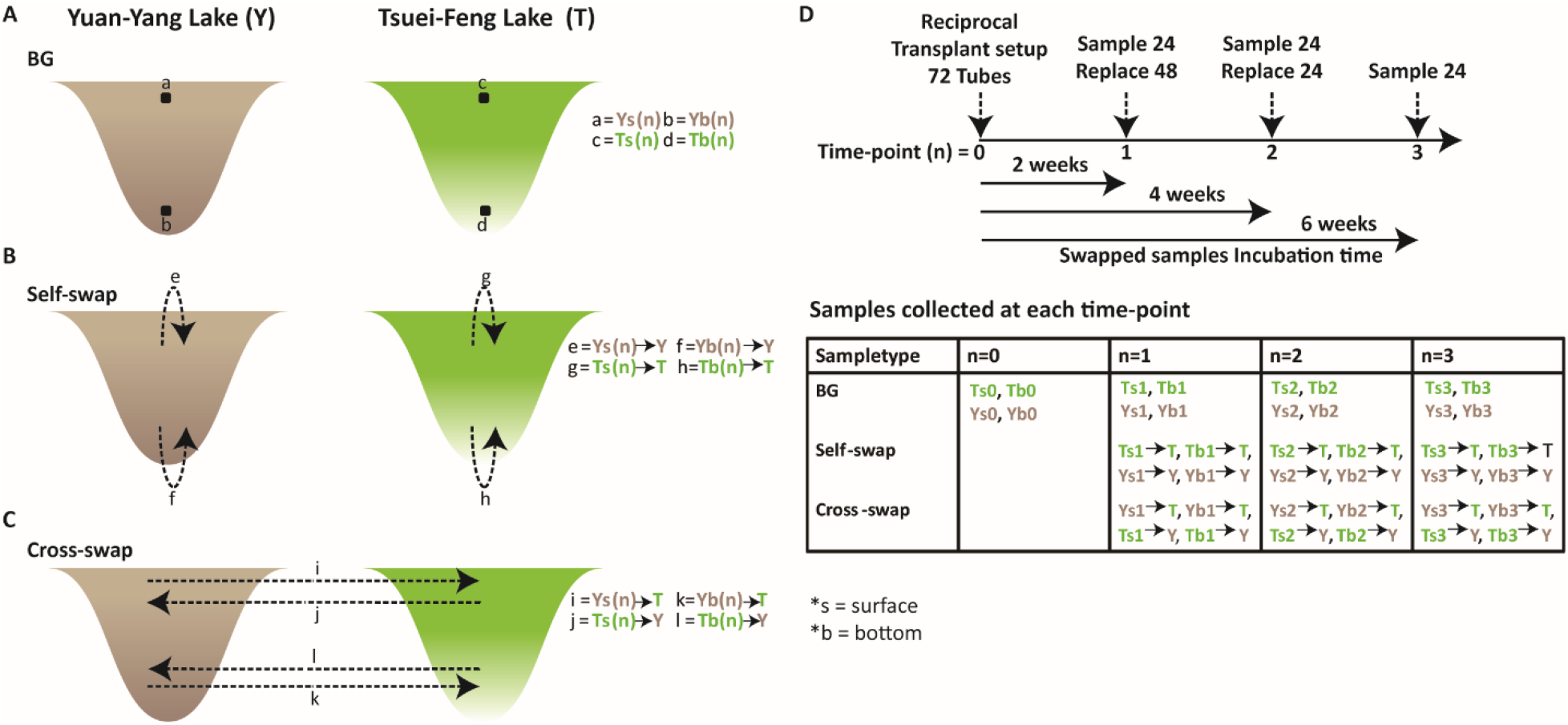
Experimental setup and sampling strategy. **(A–C)** Schematic representation of the background (BG) and reciprocal transplant (self-swap and cross-swap) samples; small letters from a-l denote samples from each group (BG, self-swap, and cross-swap). (**D**) Sampling strategy: water samples were swapped (self-swap and cross-swap) at time point n=0 and incubated; samples were collected every 2 weeks (BG, self-swap, and cross-swap) and the remaining sampling tubes were replaced with fresh tubes to avoid algal overgrowth.

Seventy-two dialysis tubes (36 surface and 36 bottom) filled with water samples (~500 mL each) were incubated for 6 weeks after swapping (time point (n)=0) (Fig. 1D, Supplementary Fig. 1A). Every two weeks, samples from 24 tubes (12 from each lake (6 self-swap and 6 cross-swap; surface and bottom)) along with BG control samples at each time point were collected and the remaining dialysis tubes containing swapped water samples were replaced with new tubes to avoid algal growth on their outer surface (Fig. 1D, Supplementary Fig. 1A). Collected water samples from tubes were filtered using a piece of gauze (to remove large debris, e.g. leaves), then sand and dust particles were removed using an 11-μm filter. Finally, 1.5 L of water was filtered using a 0.22 μm filter (diameter 0.47 mm, Advantec); ~200 mL and ~1 L was stored at −20°C until amplicon-based nucleic acid extraction and metagenome DNA extraction, respectively. Three replicates (n=3) of each sample were collected for amplicon sequencing and no replication was performed for whole-metagenome sequencing.

### Nucleic acid extraction, PCR amplification, and sequencing

Total genomic DNA from filtered water samples was extracted using the UltraClean Soil DNA Kit (MoBio, Solana Beach, CA, USA) following the manufacturer’s protocol. RNA from water samples was extracted using the UltraClean Soil RNA Kit (MoBio, Solana Beach, CA, USA), and residual DNA was removed using a TURBO DNA-Free Kit (Invitrogen, Carlsbad, CA, USA). The nucleic acid yield was determined by NanoDrop ND-1000 UV-Vis Spectrophotometer (NanoDrop Technologies, Inc., Wilmington, DE, USA).

Complementary DNA (cDNA) was synthesized from purified RNA using the SuperScript IV First-Strand Synthesis System for RT-qPCR (Invitrogen, Carlsbad, CA, USA) following the manufacturer’s protocol. To prepare 16S rRNA gene-transcripts, PCR amplification was conducted to target the V6–V8 hypervariable regions of the bacterial 16S rRNA gene using the bacterial universal primers −968F (5’AACGCGAAGAACCTTAC-3’)^37^ and 1391R (5’-ACGGGCGGTGWGTRC-3’)^38^. The reaction mixture contained 1 μL of 5 U TaKaRa *Ex Taq* HS (Takara Bio, Otsu, Japan), 5 μL of 10× *Ex Taq* buffer, 4 μL of 2.5 mM deoxynucleotide triphosphate mixture, 1 μL of each primer (10 μM), and 1–5 μL (10–20 ng) of template DNA for a total volume of 50 μL. PCR was performed under the following conditions: 94°C for 5 min; 30 cycles of 94°C for 30 s, 52°C for 20 s, and 72°C for 45 s; and a final extension of 72°C for 10 min.

DNA-tagging PCR was used to tag each PCR product of the bacterial 16S rRNA gene’s V6–V8 region^37^. The tag primer was designed with four overhanging nucleotides; this arrangement yielded 256 distinct tags—at the 5′ end of the 968F and 1391R primers—for bacterial DNA. The tagging PCR conditions comprised an initial step of 94°C for 3 min; 5 cycles at 94°C for 20 s, 60°C for 15 s, 72°C for 20 s; and a final step at 72°C for 2 min.

### Amplicon and metagenome sequencing

Illumina sequencing (MiSeq) was performed on pooled 40-ng lots of uniquely marked samples by Yourgene Biosciences, Taiwan. Six TruSeq DNA PCR-Free libraries (three for 16S rRNA gene and three for 16S rRNA gene-transcripts) were prepared for 2×300 bp paired-end reads by Yourgene Biosciences. Raw reads were sorted and primers were removed before further analyses. Amplicon data—i.e. 16S rRNA gene and 16S rRNA gene-transcripts—were used for total and active microbial community composition analysis, respectively (Supplementary Fig. 1B).

For whole-metagenome sequencing, the total genomic DNA was extracted using the UltraClean Soil DNA Kit (MoBio, Solana Beach, CA, USA) following the manufacturer’s protocols. Extracted DNAs from the 39 samples (Supplementary Fig. 1B) were sent to Yourgene Biosciences (Taipei, Taiwan) for library preparation without amplification (PCR-Free library) and sequencing by the Illumina MiSeq system (USA) with a read length of 2 × 300 bp. Whole-metagenome samples were used to determine the functional profile of the microbial community (Supplementary Fig. 1B).

### Community level physiological profiling

The carbon assimilation potential of the microbial communities in the two lakes was assessed with Biolog EcoPlates^TM^ (Biolog Inc., Hayward, CA, USA), hereafter written as ecoplates. Undiluted water (150 μL) from each sample was added to the ecoplate wells, each containing 31 different carbon sources (and one control well with no carbon source). Plates were incubated at 25°C and the absorbance of each well was measured at 590 nm using an automatic optical density microplate reader (Spectramax M2, Molecular Devices, San Jose, CA, USA) every 24 hours for 7 days. On day 7, absorbance values from the ecoplates were used to calculate the average well color development (AWCD) in the plate, as expressed by the equation below.

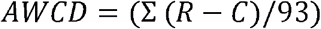

where C and R represent the absorbance value in the blank well and substrate, respectively. Negative values were converted to 0. The absorbance values were then treated and normalized using AWCD. A substrate with a normalized absorbance value >1 was considered to be metabolized by the microbial community.

### Sequencing data analyses

#### Amplicon data analyses

Paired-end 16S rRNA gene and 16S rRNA gene-transcript reads were sorted into samples using an in-house script. Sequences in each sample were screened by MOTHUR v1.3.81^39^ to keep those 1) of lengths >390 bp and <450 bp, 2) with 0 ambiguous bases, and 3) with homopolymers <8 bp. Chimeric reads were detected and removed using UCHIME^40^ by USEARCH v11^41^. Qualified sequences were retained for further analysis. Qualified and non-chimeric reads were analyzed with UNOISE3^42^ to obtain zero-radius operational taxonomic units (zOTUs), which are equivalent to exact sequence variants. zOTUs were classified with taxonomic labels using a combination of two databases—the generic 16S rRNA gene database SILVA132^43,44^ and the custom freshwater database TaxASS^45^—to obtain a fine-scale taxonomic classification in Mothur on a per-sample basis with a pseudo-bootstrap cutoff of 80%.

#### Whole-metagenome data analyses

Metagenomic reads (2 × 300 bp) obtained from the Illumina MiSeq system were quality-checked and adapters removed using Cutadapt v1.4.2^46^, quality trimming of reads was performed with Seqtk v. 1.2. −r94 (phred score <20, short read cut off <35 bp)^47^, and singlet reads were discarded. Quality filtered paired reads were merged using FLASH version 1.2.11^48^. These merged reads were searched for similarity using DIAMOND^49^ (e-value cut off 1e-5) against the Clusters of Orthologous Groups database (COGs) (last downloaded 2019) at the COG-Class and -Family levels to obtain functional profiles of metagenome samples. COG-Family represented in 90% of the samples with at least 10 reads each were used for the downstream analysis.

### Statistical analyses

#### Similarity

All statistical analyses and plots were conducted in R (R Core Team, 2018). Alpha diversities (Richness, Shannon, Chao1, and Inverse Simpson (Inv. Simpson)) were calculated after rarifying the samples to an even depth of 23,616 reads using the estimate_richness function from phyloseq^50^, with subsampling libraries with replacement 100 times and averaging the diversity estimates from each trial. Abundance profiles from the 16S rRNA gene and 16S rRNA gene-transcript amplicon datasets were processed with the R packages phyloseq, vegan^51^, and ggplot2^52^ for downstream analysis and visualization. All the abundance profiles were log (x+1)-transformed to achieve normality. Principal coordinate (PCoA) ordination analysis was performed to compare community compositions as well as functional attributes using the Bray-Curtis distance metric unless otherwise stated. 16S rRNA gene and gene-transcripts datasets were compared using Procrustes and Permutest analyses from the vegan package.

#### Significance

Kruskal-Wallis test followed by pairwise Wilcox test was used to statistically test for significant differences in alpha diversity estimates between groups. Procrustes and PROTEST analyses (999 permutations) was used to infer congruence between 16S rRNA gene and 16S rRNA gene-transcript datasets. Permutational Analysis of Multivariate Dispersion (PERMDISP) was used to test for homogeneity of multivariate dispersion between groups, followed by Permutational multivariate analyses of variance (PERMANOVA) to statistically test for differences in community compositions and functional attributes between groups. PERMANOVA and PERMDISP analyses were performed using the functions Adonis and betadisper, respectively, in the vegan package. Welch’s *t*-test implemented in STAMP^53^ was used to test (Bonferroni corrected *p*<0.05) differentially abundant COG-Class categories between groups. The relationship between community and functional attributes was statistically tested with regression analysis and ANOVA with the stats package in R. Functional redundancy was quantified using a custom script written in R to calculate *L*-divergence^54^. The effect of the transplant experiment on carbon assimilation via ecoplates analysis was tested using an unpaired Student’s *t*-test (significance: *p*<0.05).

### Data availability

All the sequencing data were deposited into the NCBI database under the BioProject PRJNA596577.

## Results

### zOTU filtering

In total, we observed 926 zOTUs for the total community (16S rRNA gene), eight of which were classified as Archaea, two as Eukaryota, 45 as chloroplast, two as mitochondria, and seven as unknown. After removing 54 zOTUs, 851 and 858 bacterial zOTUs remained from surface and bottom water samples, respectively. A total of 1044 zOTUs were identified in the 16S rRNA gene-transcript dataset (active community), of which 54 zOTUs were chloroplast, 21 were mitochondria, and four were unknown. After removing 79 zOTUs, the surface and bottom samples had 945 and 957 zOTUs, respectively. Bacterial zOTUs from the total and active community datasets were used for all downstream analyses.

### Disturbance affects bacterial diversity and richness

Alpha diversity estimates of BG control samples (Ys *vs* Ts and Yb *vs* Tb) from the total community identified that Ys and Yb had higher Richness, Shannon, Inv. Simpson, and Chao1 estimate compared to Ts and Tb samples, respectively (Supplementary Fig. 2A and 2B). None of the diversity estimates were significantly different for surface samples, but Shannon and Inv. Simpson diversity estimates were significantly different for bottom samples (*p*<0.05). When cross-swapped samples were compared to their self-swap counterparts, we found that all calculated diversity estimates decreased when Y lake’s surface and bottom samples were swapped with T lake’s surface and bottom and compared with their self-swap counterparts (Ys→T *vs* Ys→Y and Yb→T *vs* Yb→Y), respectively (Supplementary Fig. 2A and 2B). No significant differences were identified between the compared cross and self-swap groups. However, for the other way round (Ts→Y *vs* Ts→T and Tb→Y and Tb→T), all diversity estimates increased, but none significantly (Supplementary Fig. 2A and 2B).

16S rRNA gene-transcript-based alpha diversity estimates showed a similar trend, with the Inv. Simpson diversity estimate being significantly different (*p*<0.05) between Ys and Ts samples (Supplementary Fig. 3A). The richness and Chao1 were also significantly (*p*<0.05) different between Yb and Tb samples (Supplementary Fig. 3B). The cross-swap samples, however, were not significantly different in their estimated alpha diversity measures (Supplementary Fig. 3A and 3B).

### Microbial community composition of Tsuei-Feng and Yuan-Yang Lakes’ surface and bottom water

The total bacterial community composition based on the 16S rRNA gene dataset showed that the bacterial communities in the surface and bottom waters of both lakes were dominated by different bacterial classes. Ts samples were dominated by *Actinobacteria* (average relative abundance: 43.02%) and *Gammaproteobacteria* (20.69%) (Silva132 release merged *Betaproteobacteria* with *Gammaproteobacteria)*. Ys samples, on the other hand, were dominated by *Gammaproteobacteria* (60.93%) and *Bacteroidia* (16.11%). *Verrucomicrobia* was the most abundant class in both Tb (47.022%) and Yb (36.90%) samples. The class *Cyanobacteria* was only abundant in Ts samples (3.76%). In swapped samples, abundant bacterial classes remained the same in both the self-swap (Ts(n)→T, Tb(n)→T; Ys(n)→Y, Yb(n)→Y) and cross-swap (Ys(n)→T, Yb(n)→T; Ts(n)→Y, Tb(n)→Y), with changes occurring only in their relative abundances (Supplementary Fig. 4A and 4B).

The 16S rRNA gene transcript-based active bacterial community of both Ts and Tb samples were dominated by the classes *Actinobacteria* (surface: 25.51%; bottom: 28.25%)*, Bacteroidia* (25.02%; 27.82%), *Cyanobacteria* (20.49%; 16.16%), and *Alphaproteobacteria* (18.74%; 17.50%), whereas Ys and Yb samples were dominated by *Gammaproteobacteria* (41.07%; 40.38%)*, Alphaproteobacteria* (20.28%; 9.82%) and *Bacteroidia* (27.96%; 27.45%) (Fig. 2A and 2B)*. Cyanobacteria* were only abundant in the Tsuei-Feng samples. Swapped samples had the same most abundant classes, with changes only occurring in the relative abundance profiles. *Actinobacteria* relative abundance decreased in self-swap samples (Ts→T (7.34%) and Tb→T (4.76%)), *Cyanobacteria* relative abundance decreased in Ts(2)→Y (5.52%), before increasing again in Ts(3)→Y (28.38%) and also in Tb(2)→Y (2.47%) and Tb(3)→Y (2.97%) samples. *Alphaproteobacteria* relative abundance increased in cross-swap samples (Ys(1)→T, Ys(2)→T, and Ys(3)→T) and *Fimbriimonadia* abundance increased in only Ys(3)→T (10.91%) and Yb(3)→T (2.86%) samples compared to their self-swap counterparts (Ys(3)→Y (0.02%) and Yb(3)→Y (0.02%)) (Fig. 2A and 2B).

**Fig 2.**
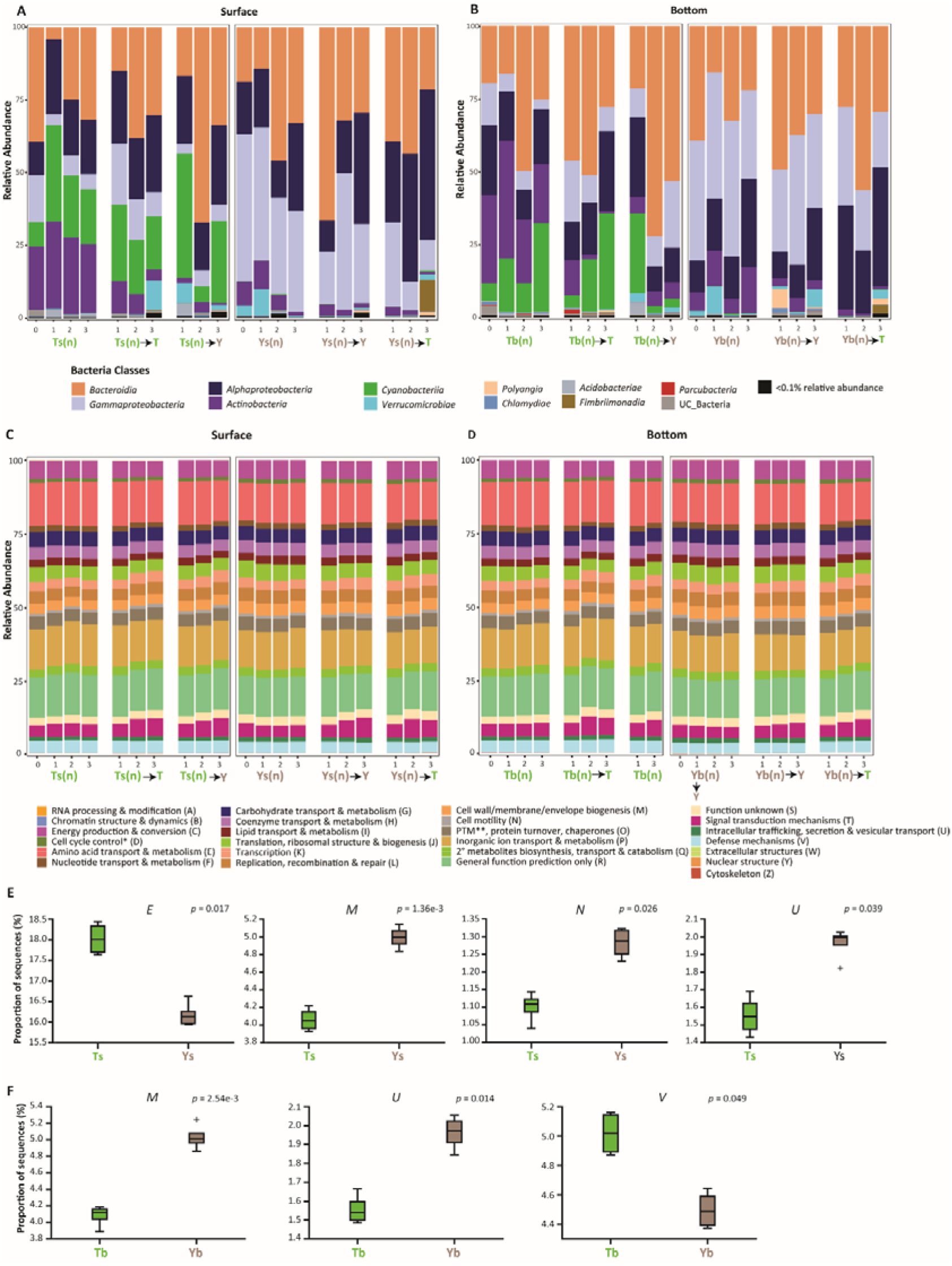
Tsuei-Feng and Yuan-Yang Lakes’ surface and bottom waters showed different bacterial communities but similar functional profiles. **(A–B)** Bacterial community profiles at the class level for the both lakes surface **(A)** and bottom (**B)** waters were diverse and did not show significant changes after cross-swapping. (**C–D)** Functional profile at the COG-Class level; the functional profiles appeared stable and very similar across all sampling groups (BG, self-swap, and cross-swap) in surface and bottom waters. (**E–F)** Statistical analysis identified significantly different (Welch’s t-test, Bonferroni corrected *p*<0.05) COG-Classes between the background (BG) surface and bottom sampling groups between the two lakes; no other groups showed significant difference. COG-Classes E, M, N, U, and V are the same as shown in (C–D)

### Microbial functional profiles of Tsuei-Feng and Yuan-Yang Lakes’ surface and bottom waters

The whole-metagenome-based functional profiles derived at the COG-Class level appeared very similar and stable across sample types (BG, self-swap, and cross-swap) and time for both surface and bottom waters (Fig. 2C and 2D). COG-Class categories, Amino acid transport and metabolism (E), Inorganic ion transport and metabolism (P), Energy production and conversion (C), and General functional prediction (R) only accounted for ~50% of the total annotated functional abundance. Despite appearing very similar and stable, the statistical comparison of the surface and bottom water samples from both lakes showed that COG-Classes E (Welch’s *t*-test, *p.adj=*0.017) and Defense mechanism (V) (Welch’s *t*-test, *p.adj=*0.049) had a significantly high proportion of reads in the BG control samples (Ts and Tb) (Fig. 2E and 2F). Cell wall/membrane/envelope biogenesis (M) (only surface samples) (Welch’s *t*-test, *p.adj=*1.36e-3), cell motility (N) (surface: Welch’s *t*-test, *p.adj=*0.026; bottom: Welch’s *t*-test, *p.adj=*2.54e-3), and Intra-cellular trafficking, secretion, and vesicular transport (U) (surface: Welch’s *t*-test, *p.adj=*1.36e-3; bottom: Welch’s *t*-test, *p.adj=*0.014) had significantly higher proportions of reads annotated in Ys and Yb samples compared to Ts and Tb, respectively (Fig. 2E and 2F). No significant differences were observed in any other comparisons.

### Establishing community functional relationship

We conducted PCoA analysis to compare the active bacterial community composition at the finest resolution for zOTUs between the two lakes and understand the dynamics of microbial community following the reciprocal transplant. Community composition was significantly different between lakes at the zOTU level, with the first two axes of PCoA explaining a total of 62% and 58.8% of the variance in surface and bottom samples, respectively (Fig. 3A and 3C). Moreover, the inoculum was the most significant factor driving this difference in community composition between the surface (Adonis _(inoculum)_: F_1, 19_ = 20.58, R^2^ = 0.50, *p* = 0.001) and bottom waters (Adonis _(inoculum)_: F_1,19_ = 18.11, R^2^=0.46, *p*=0.001) of the two lakes; no significant difference was identified for community dispersion (PERMDISP_(inoculum)=_ *p*>0.05; PERMDISP_(incubating-lake)=_ *p*>0.05) in surface and bottom waters. Incubation in the lake had a weak effect on the surface water community (Adonis_(incubating-lake):_ F_1,19_ = 2.48, R^2^ = 0.060, *p* = 0.047) compared to the inoculum. Samples from the lakes—irrespective of their type (BG, self, and cross-swap)—clustered distinctly (Fig. 3A and 3C). Notably, swapped samples showed slight variation (Fig. 3A and 3C, denoted by dotted grey arrows) compared to BG control samples. Overall, the active (Fig. 3A and 3C) and total (Supplementary Fig. 4C and 4D) bacterial communities appeared resistant to change, even when swapped and incubated in a different environment. Procrustes analysis revealed congruence between total and active bacterial community compositions for surface (PROTEST: m_12_^2^: 0.0581; correlation: 0.9705; *p = 0.001*) and bottom (PROTEST: m_12_^2^: 0.0518; correlation: 0.960; *p = 0.001*) samples. Hence, we used only the active bacterial community dataset to quantify the community functional relationships at a later stage.

**Fig 3.**
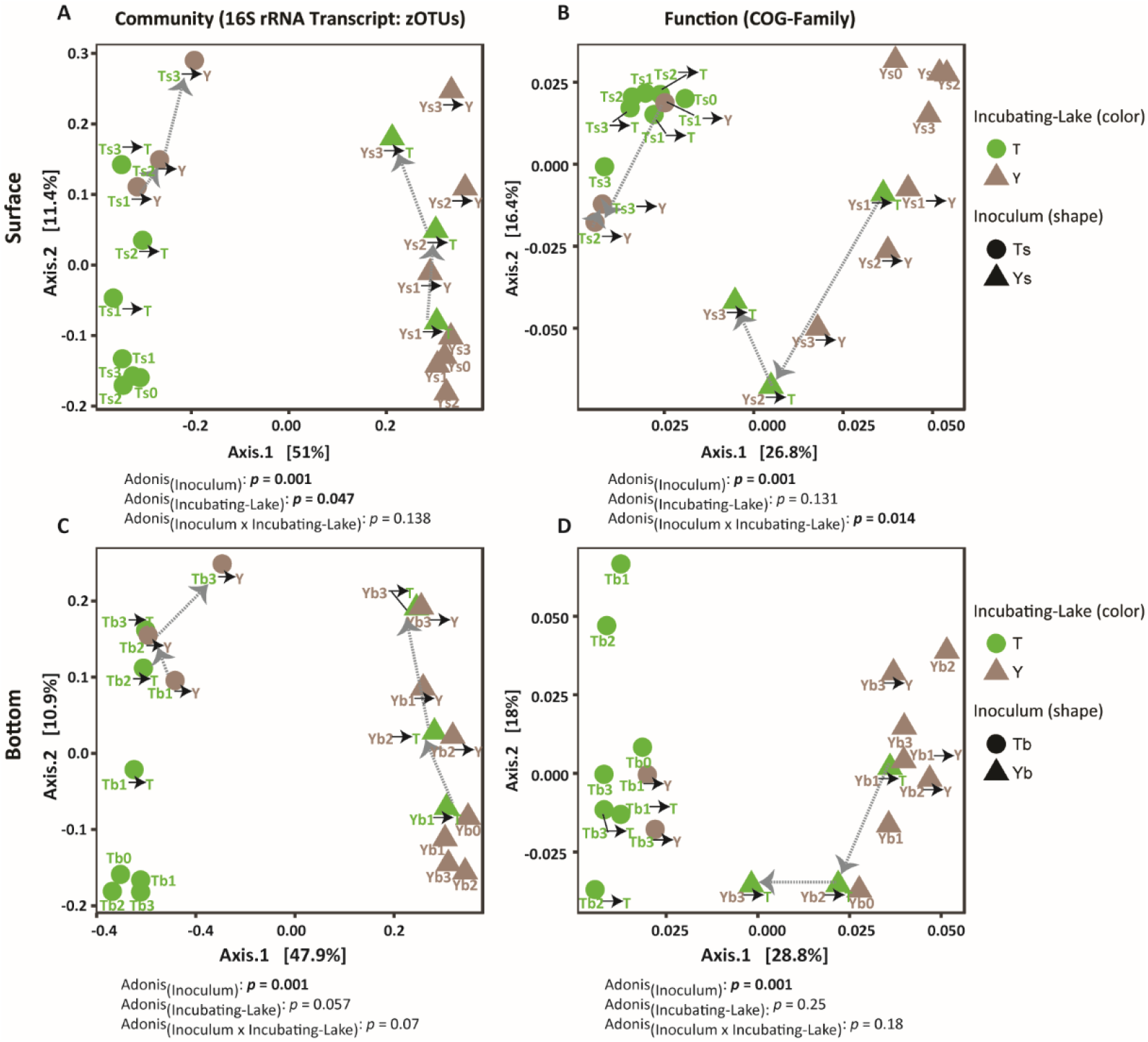
Principal coordinate analysis of bacterial communities and functional profiles. Active bacterial community compositions (based on zero-radius Operational Taxonomic Units (zOTUs)) from the Bray-Curtis distance metric differed most between lake surface **(A)** and bottom **(C)** waters, even after a reciprocal transplantation, with the inoculum being a significant factor (*p*<0.05). COG-Family-based functional profiles found differences between surface **(B)** and bottom **(D)** waters; the inoculum and inoculum x incubating lake were significant factors for the surface, whereas the inoculum alone was the significant factor for bottom waters. In addition, cross-swap samples tended to shift away from their self-swap counterparts for function, whereas in the case of community compositions, they shifted along with their self-swap counterparts (marked by dotted grey arrows). Functional profiles were obtained from the metagenomes after COG-Family classification at the read level.

COG-Class provides a general view of functional characterization, so we used COG-Family level (finest metabolic resolution possible) to delineate the effect of reciprocal transplant on the microbial functional profile. The PCoA analysis at the COG-Family level showed that the functional profiles of the two lakes was more similar than were their community compositions (Fig. 3B and 3D). The first two PCoA axes explained 43.2% and 36.8% of the variance in the surface and bottom samples, respectively. Inoculum was still the most significant factor driving the observed differences between functional profiles of the two lakes in surface (Adonis _(inoculum)_: F_1,19_ = 6.43, R^2^ = 0.24, *p* = 0.001) (Fig. 3B) and bottom (Adonis_(inoculum)_: F_1,18_ = 6.44, R^2^ = 0.26, *p=0.001*) (Fig. 3D) samples, though this variable was relatively weaker when used to compare community compositions. Interestingly, we observed shifts in cross-swap (surface: Ys(1)→T, Ys(2)→T, Ys(3)→T, Ts(1)→Y, Ts(2)→Y, Ts(3)→Y) samples away from their self-swap counterparts and towards the incubating lake. These shifts were more apparent for Ys(n)→T, Ts(n)→Y, and Yb(n)→T samples than for Tb(n)→Y, potentially because the Yb(2)→T sample was lost during processing (Fig. 3B and 3D, denoted by arrows). Furthermore, for surface samples, we observed that the interaction term *inoculum* x *Incubating-Lake* (Adonis_(inoculum x_ Incubating-Lake: F_1,19_ = 2.30, R^2^ = 0.08, *p* = 0.014) also had a significant but weak effect that was not observed in lake bottom water samples (Fig. 3B and 3D). PERMDISP did not reveal any significant difference (*p*>0.05) for the inoculum and incubating lake for surface and bottom waters. Shifts in microbial functional profiles, when incubated in different environments and under the influence of both the inoculum and incubating lake, shed light on the ability of microorganisms to adapt to their local environment.

### Quantifying community functional relationships

To quantify the microbial community functional relationships, we performed a linear regression analysis using Bray-Curtis similarity (1-Bray-Curtis distance) on the community compositions and functional profiles. We observed a significant and positive linear relationship for the surface (Adj. R^2^ = 0.5065, *p*<0.00001) and bottom (Adj R^2^ = 0.4592, *p*<0.00001) (Fig. 4A and 4B) levels. Interestingly, the range of similarity between the community composition of lakes varied from <20% to >70% (x-axis, Fig. 4A and 4B), whereas their functional attributes had higher similarities and a narrower range of <88% to >94% (y-axis, Fig. 4A and 4B). Furthermore, we used *L*-divergence to quantify the community functional relationships. *L*-divergence can range from 0 (identical) to 2 (highly divergent). Results showed high *L*-divergences for community compositions, with a mean±s.d. of 1.21±0.54 for the surface (Fig. 4C) and 1.17±0.51 for bottom (Fig. 4E), and a low *L*-divergence of 0.04±0.01 for the surface (Fig. 4D) and bottom (Fig. 4F) functional attributes. A positive linear relationship, with low community similarity but a higher similarity in functional attributes, points to the presence of partial functional redundancy. Furthermore, a highly divergent community but a relatively low divergent metabolic functional profile provides further support this finding of functional redundancy.

**Fig 4.**
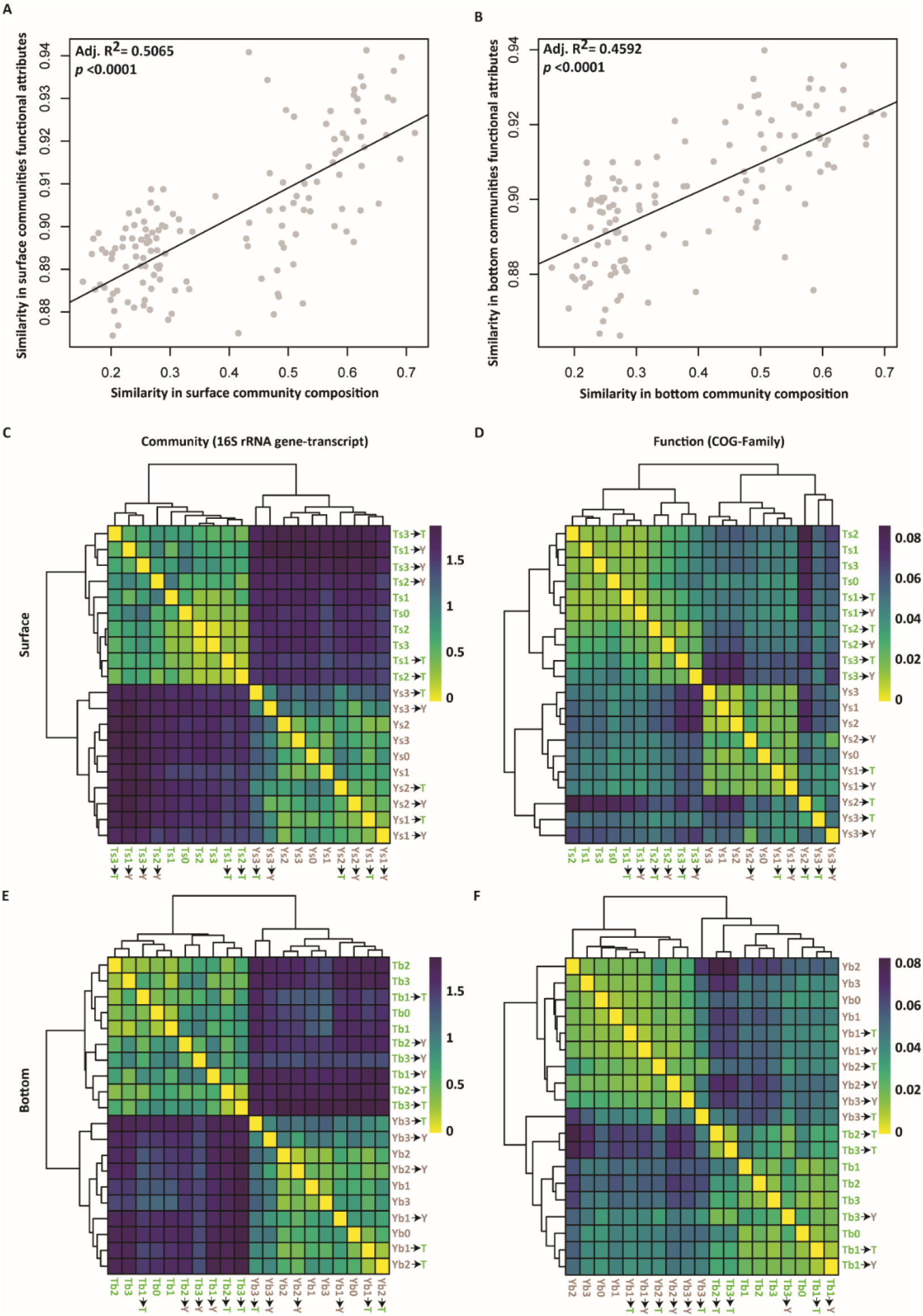
The similarities in community composition are related to similarities in functional attributes, but at a different range. Regression analysis identified a significant positive linear relationship between the similarity in community composition and overall functional attributes with a different similarity range (Bray-Curtis similarity, Community: >20–<70%; Function: >88–<94%) for surface **(A)** (R^2^ = 0.5065, *p*<0.0001, *F* test) and bottom waters **(B)** (R^2^ = 0.4592, *p*<0.0001, *F* test). The similarity (1-Bray Curtis dissimilarity) in functional attributes was measured by directly comparing COG-Family profiles for each surface and bottom metagenome samples and similarity between communities was estimated from the composition of 16S rRNA gene transcripts dataset. Community compositions were highly divergent between the two lakes in both the surface **(C)** (*L*-divergence=1.21±0.54) and bottom samples **(E)** (*L*-divergence=1.17±0.51), but functional profiles were less divergent (*L*-divergence=0.04±0.01). *L*-divergence was calculated for the community using the 16S rRNA gene-transcript dataset at zOTU levels and using a metagenomes dataset after COG-Family classification at the reads level for the surface and bottom separately.

### Microbial metabolic activity is influenced by disturbance

We used ecoplates as a proxy to measure the metabolic activity of the microbial community in the two lakes and monitor changes after the reciprocal transplant experiment. Similar substrates were utilized by the surface and bottom microbial communities present in both lakes (Supplementary Figs. 5 and 6). The results indicated a widespread phylogenetic diversity in carbon metabolism.

BG samples of Y lake (Ys and Yb) had higher average well color development (AWCD) compared to those of T lake (Ts and Tb) (Supplementary Fig. 7A and 7B). After the reciprocal transplantation and incubation for 2 weeks, we observed that cross-swap samples Ys→T and Yb→T had significantly (*p*<0.05) lower AWCD compared to their self-swap counterparts (Ys(1)→T vs Ys(1)→Y; Yb(1)→T vs Yb(1)→Y) (Supplementary Fig. 8A and 8B), whereas Ts→Y and Tb→Y had significantly (*p*<0.05) higher AWCD compared to their self-swap counterparts (Ts→T and Tb→T) (Supplementary Fig. 8C and 8D). Subsequently, after 4 weeks of incubation, we observed that the AWCD of cross-swap bottom samples (Yb(2)→T and Tb(2)→Y) was not significantly (*p* >0.05) different to that of their self-swap counterparts (Yb2→Y and Tb2→T) (Supplementary Fig. 8B and 8D). However, surface samples (Ys(2)→T and Ts(2)→Y) had significantly different (*p*<0.05) AWCD values compared to their self-swap counterparts (Ys(2)→Y and Ts→T) (Supplementary Fig. 8A and 8C). Finally, after 6 weeks of incubation in lakes, we observed that cross-swap sample Ts(3)→Y continued to have significantly (*p*<0.05) higher AWCD compared to its self-swap counterpart Ts(3)→T (Supplementary Fig. 8C). But contrasting results were obtained for Ys(3)→T *vs* Ys(3)→Y: Ys(3)→T had significantly (*p*<0.05) higher AWCD compared to Ys(3)→Y (Supplementary Fig. 8A). In addition, in the case of bottom samples, Tb(3)→Y *vs* Tb(3)→T did not have significantly (*p* >0.05) different AWCD values, but Yb(3)→T had significantly (*p*<0.05) higher AWCD compared to Yb(3)→Y (Supplementary Fig. 8B and 8D).

Shifts in the microbial metabolic activity, measured using AWCD as a proxy, in cross-swap samples after incubation in a contrasting environment indirectly points to the influence of the local environment in shaping the rate of substrate utilization. However, while it is not direct *in situ* evidence, our analysis does offer evidence that shifts in metagenome-derived functional profiles after water samples were reciprocally transplanted in different lakes.

## Discussion

Our understanding of community functional relationships and their magnitudes in natural environments remains poor owing to a lack of *in situ* empirical time-series studies. In this setting, the main motivation of our study was to empirically test the community functional relationships in the aquatic ecosystem with an *in situ* reciprocal transplant experiment. We found that the two lakes have contrasting microbial communities that appear to be resistant to change, even when swapped and incubated in a different environment. Community functional attributes tend to become more similar to the functional profile of the incubating lake. We also report the presence of partial functional redundancy in the aquatic microbiome, with high divergence in the community compared to the community’s functional attributes. To the best of our knowledge, this is the first study to 1) examine the function redundancy hypothesis in surface and bottom lake waters with a time series *in situ* reciprocal transplant experiment and 2) make use of amplicon and metagenome sequencing complemented with an ecoplate-based functional assay to establish community functional relationship.

### Bacterial diversity and composition in the lakes

A small volume of freshwater can contain thousands of dissolved organic molecules with varying complexities^55^; this chemical diversity has been known to positively influence microbial diversity and ecosystem functioning^56^. The bacterial communities identified in the two studied lakes are typical of freshwater communities, which are often dominated by *Actinobacteria, Alpha*- and *Gamma-proteobacteria* (now including *Betaproteobacteria*)*, Bacteroidia, Cyanobacteria,* and *Verrucomicrobia*^23,57–60^. Both lakes contained generalist taxa, though at different abundances (Fig. 2A and 8B and Supplementary Fig. 4A and 4B), and this highlights the influence of deterministic (local environment) and stochastic processes (dispersion)^61,62^. Recent studies have also pointed out that deterministic and stochastic processes have the potential to influence the structure and function of microbial communities at different scales^63^. Furthermore, we observed that microbial community diversity is influenced by swapping (disturbance), including self-swapping and cross-swapping (Supplementary Figs. S2 and S3). Microbial community diversity is known to be associated with lake mixing regimes, a natural disturbance to bacterial communities^64^.

### Community composition appeared resistant to disturbance after transplantation

The total and active bacterial community compositions of BG control samples were significantly different between the lakes. When comparing transplant samples, we found that cross-swap samples clustered with their self-swap counterparts (Fig. 3A and 3C and Supplementary Fig. 4C and 4D), even after incubation in a contrasting environment for six weeks. Small deviations were observed in swapped samples over time when compared to BG control samples, and we believe this can be attributed to the effect of tubing on the microbial community. Furthermore, only inoculum had a significant effect (surface: Adonis_(inoculum)_ *p=*0.001; bottom: Adonis_(inoculum)_ *p=*0.001) on the clustering of samples. Our results are similar to those of Langenheder et al.^25^ and others^12,65^, who suggest that the bacterial communities in these environments are resistant to change. Despite strong evidence of resistance, we cannot rule out another property of microbial community: resilience, or the existence of an alternate stable state of a microbial community^12,65^. It is possible that the microbial community shifted soon after the transplant and then recovered to an alternate state before the next sampling (2 weeks later), as observed from deviations in the swapped samples. Bacterial communities in soil (reviewed in ^66^), lakes, and reservoirs are known to recover quickly after natural disturbances such as lake mixing^64^ and typhoons^26^. Furthermore, earlier studies have shown that microbial communities possess properties of both resilience and resistance to change^12,65,66^.

### Functional attributes and metabolic ability are influenced by the local environment

We hypothesized that swapping water from one lake to another and vice versa would have consequences for the functional attributes of both lakes’ microbial communities, and used a combination of metagenomic and ecoplate approaches to test this. A similar hypothesis was tested and supported in an earlier study^67^. At the broad COG-Class level, we observed that the community functional profile appeared very similar and stable (Fig. 2C and 2D) based on relative abundance across all groups, with only a few COG-Classes (E, M, N, U, and V) showing a significant difference (only in BG control samples) (Fig. 2E and 2F), much like community profile observed in earlier studies on different ecosystems like oceans^2^ and water reservoirs^26^. At a finer resolution (COG-Family), interesting insights were observed: cross-swap samples tended to diverge from their self-swap counterparts. These shifts were in varying degrees at the surface and bottom levels and also between the lakes (Fig. 3B and 3D), highlighting the influence of the local environment as well as the resolution of the analyses (COG-Class *vs* COG-Family), though the former was not quantified. Deterministic factor-like environmental filtering has been known to influence the functional composition of microbial communities in several ecosystems, including plants^9^ and oceans^3^.

In recent years, ecoplate-based analyses have been applied as proxies for the functional diversity of microbial communities^29,31,68^ and related community functional relationships in different ecosystems. It is important to note that all substrates in an ecoplate have a carbon backbone and that carbon metabolism is phylogenetically widespread^67^. In light of this, several studies have reported the presence of functional redundancy in ecosystems^29,69^ when using ecoplates as a proxy for microbial community functions. Since many generalists dominated the microbial community of two lakes studied here, it was no surprise that most of the ecoplate substrates were utilized (Supplementary Figs. 5 and 6), suggesting the presence of functional redundancy concerning carbon metabolism. Furthermore, microbes (especially bacteria) are known to overcome changes in the environment by expressing a range of metabolic capabilities or the property of mixotrophy^70^ and varying affinities to organic molecules. Interestingly, using AWCD values as a proxy for metabolic activity, we observed an increase in AWCD when water from T lake was swapped with that of Y lake (Supplementary Figs. 7 and 8) and a decrease in AWCD when Y lake water was swapped with T lake water (Supplementary Figs. 7 and 8) and compared with their respective self-swap counterparts. Changes in metabolic activity provide indirect evidence of the influence of the local environment on the metabolic activity of the microbial community. It is worth noting here that very similar substrates were utilized by the bacterial communities in both lakes, suggesting the presence of metabolic plasticity. Earlier studies have shed light on the metabolic plasticity that microorganisms possess allowing them to survive in lakes and maintain overall ecosystem functioning^68,71^.

### Partial functional redundancy in aquatic ecosystems

Establishing community functional relationships has been at the forefront of functional redundancy research. In recent years, many studies have reported partial functional redundancy in different ecosystems^3,5,17,25,30^. Using linear regression analysis, we observed that there is a positive correlation between the community and functional similarity in the aquatic microbiome for surface (Adj. R^2^: 0.5065, *p*<0.0001) and bottom (Adj. R^2^: 0.4592, *p*<0.0001) waters. Furthermore, we observed that functional attributes have a narrow range with higher similarity as compared to the communities’ broader range with lower similarity, suggesting the presence of partial functional redundancy (Fig. 4A and 4B). Using the *L*-divergence measure, we quantified the scale of partial functional redundancy and identified a relatively highly divergence in the community (*L*-divergence_surface_: 1.21±0.54; *L*-divergence_bottom_: 1.17±0.51) and low divergence in that community’s functional profile (*L*-divergence_surface_: 0.04±0.01; *L*-divergence_bottom_: 0.039±0.01), confirming the scale of partial functional redundancy. Our results contrast with those of Galand et al.^3^, who identified similar scales for community and functional similarity in ocean and suggested the absence of strict functional redundancy. There are two possible explanations for this: first, the differences between freshwater and marine aquatic environments, and second, the duration (the present study was performed for 6 weeks while Galand et al.^3^ spanned 3 years). Seasons are known to affect the microbial community composition of oceans and impose broad-scale changes in community composition; it is therefore possible that seasons also affect microbe-mediated processes, resulting in an absence of functional redundancy.

### Limitations of this study

In this study, we collected samples at 2-week intervals after reciprocal transplantation and hence cannot rule out the possibility that microbial communities posses resilience over these long intervals due to our sampling strategy; therefore, future studies should use much shorter sampling intervals to better address how microbial communities compensate for disturbance. Furthermore, as the scale (pulse and press) and duration of a disturbance can have significant effects on the microbial community and its functions, this attribute remains unexplored through our experimental setup. Another limitation of our study is the use of ecoplates as a proxy for metabolic activity: though this approach provides important clues about overall metabolism by measuring carbon metabolism, it only reflects the growth potential of bacterial groups able to metabolize the available substrates and grow under *in vitro* conditions. A meta-transcriptomics approach could have proven better at delineating the influence of disturbance on the microbial mediated process. We acknowledge that functional redundancy is a challenging concept and difficult to quantify, and we still do not know its presence, magnitude, and variability in different ecosystems, so care should be taken when using results from this study to make implications about other ecosystems, as local environmental aspects and ecosystem in study can be significant contributors to the outcomes obtained.

## Conclusion

Few *in situ* long term experiments have been conducted to empirically test the hypothesis of functional redundancy. For the first time, we conducted reciprocal transplants for surface and bottom waters of two lakes with disparate trophic states, and monitored the changes in microbial community compositions and functions over six weeks. We established that the microbial community remains resistant to change after being transplanted into a different environment, but the functional profile tends to become more similar to that of the new local environment. A statistically significant and positive linear relationship was observed between community composition and functional attributes, pointing to the presence of partial functional redundancy. We also noted that broad functions such as carbon metabolism support the presence of functional redundancy and that metabolic activity is influenced by the local environment. Based on our analyses, we conclude that there is partial functional redundancy in this aquatic microbiome with a highly diverse community but a very similar functional composition. This research further supports the presence of partial functional redundancy in aquatic ecosystems and underscores the importance of using multi-scale approaches to test community functional relationships.

## Acknowledgements

The authors thank Dr. Jiun-Yan Ding, Mr. Cheng-Yu Yang, Dr. Sonny T. M. Lee, Dr. Ching-Hung Tseng, and Dr. Carol Eunmi Lee for their assistance with sampling. K.T. acknowledges the Taiwan International Graduate Program (TIGP) for its fellowship. The authors also express their gratitude to the anonymous reviewers for their insightful comments and suggestions for improving the quality of the manuscript, and to Mr. Noah Last of Third Draft Editing for his English language editing. This study was supported by the Thematic Project at Academia Sinica (AS-103-TP-B15-3).

## Author Contributions

S.L.T. conceived of the study. K.T performed all the analyses, interpreted results, and wrote the manuscript. M.W. and C.Y. performed sampling and experiments. S.Y played an active role in the discussion and writing the manuscript. B.B., C.C., and J.T. helped with sampling and provided logistical support for the experimental setup. C.S.N. provided critical comments on the manuscript. All authors have read and approved the current version of the manuscript.

## References

1. Tringe, S. G. et al. Comparative Metagenomics of Microbial Communities. 308, 5 (2005).

2. Louca, S., Parfrey, L. W. & Doebeli, M. Decoupling function and taxonomy in the global ocean microbiome. Science 353, 1272 (2016).

3. Galand, P. E., Pereira, O., Hochart, C., Auguet, J. C. & Debroas, D. A strong link between marine microbial community composition and function challenges the idea of functional redundancy. ISME J 12, 2470–2478 (2018).

4. Gasol, J. M. et al. A transplant experiment to identify the factors controlling bacterial abundance, activity, production, and community composition in a eutrophic canyon-shaped reservoir. Limnol. Oceanogr. 47, 62–77 (2002).

5. Langenheder, S., Lindström, E. S. & Tranvik, L. J. Weak coupling between community composition and functioning of aquatic bacteria. Limnol. Oceanogr. 50, 957–967 (2005).

6. Griffiths, B. S. et al. The Relationship between Microbial Community Structure and Functional Stability, Tested Experimentally in an Upland Pasture Soil. Microbial Ecology 47, 104–113 (2004).

7. Bahram, M. et al. Structure and function of the global topsoil microbiome. Nature 560, 233–237 (2018).

8. Reich, P. B. et al. Impacts of Biodiversity Loss Escalate Through Time as Redundancy Fades. Science 336, 589–592 (2012).

9. Louca, S. et al. Functional structure of the bromeliad tank microbiome is strongly shaped by local geochemical conditions. Environmental Microbiology 19, 3132–3151 (2017).

10. Turnbaugh, P. J. et al. A core gut microbiome in obese and lean twins. 16 (2009).

11. Moya, A. & Ferrer, M. Functional Redundancy-Induced Stability of Gut Microbiota Subjected to Disturbance. Trends in Microbiology 24, 402–413 (2016).

12. Allison, S. D. & Martiny, J. B. H. Resistance, resilience, and redundancy in microbial communities. Proc Natl Acad Sci USA 105, 11512 (2008).

13. Yin, B., Crowley, D., Sparovek, G., Melo, W. J. D. & Borneman, J. Bacterial Functional Redundancy along a Soil Reclamation Gradient. APPL. ENVIRON. MICROBIOL. 66, 5 (2000).

14. Fuhrman, J. A., Cram, J. A. & Needham, D. M. Marine microbial community dynamics and their ecological interpretation. Nature Reviews Microbiology 13, 133–146 (2015).

15. Jurburg, S. D. & Salles, J. F. Functional Redundancy and Ecosystem Function — The Soil Microbiota as a Case Study. in Biodiversity in Ecosystems - Linking Structure and Function (eds. Lo, Y.-H., Blanco, J. A. & Roy, S.) (InTech, 2015). doi:10.5772/58981.

16. Strickland, M. S., Lauber, C., Fierer, N. & Bradford, M. A. Testing the functional significance of microbial community composition. Ecology 90, 441–451 (2009).

17. Fierer, N. et al. Reconstructing the Microbial Diversity and Function of Pre-Agricultural Tallgrass Prairie Soils in the United States. Science 342, 621–624 (2013).

18. Shade, A. et al. Interannual dynamics and phenology of bacterial communities in a eutrophic lake. Limnol. Oceanogr. 52, 487–494 (2007).

19. Jones, S. E., Newton, R. J. & McMahon, K. D. Evidence for structuring of bacterial community composition by organic carbon source in temperate lakes. Environmental Microbiology 11, 2463–2472 (2009).

20. Nelson, C. E. Phenology of high-elevation pelagic bacteria: the roles of meteorologic variability, catchment inputs and thermal stratification in structuring communities. ISME J 3, 13–30 (2009).

21. Judd, K. E., Crump, B. C. & Kling, G. W. Variation in dissolved organic matter controls bacterial production and community composition. Ecology 87, 2068–2079 (2006).

22. Kritzberg, E. S., Langenheder, S. & Lindstrom, E. S. Influence of dissolved organic matter source on lake bacterioplankton structure and function – implications for seasonal dynamics of community composition: Influence of DOC source on bacterioplankton. FEMS Microbiology Ecology 56, 406–417 (2006).

23. Newton, R. J., Jones, S. E., Eiler, A., McMahon, K. D. & Bertilsson, S. A Guide to the Natural History of Freshwater Lake Bacteria. Microbiology and Molecular Biology Reviews 75, 14–49 (2011).

24. Fernandez, A. S. et al. Flexible Community Structure Correlates with Stable Community Function in Methanogenic Bioreactor Communities Perturbed by Glucose. Appl. Environ. Microbiol. 66, 4058–4067 (2000).

25. Langenheder, S., Lindstrom, E. S. & Tranvik, L. J. Structure and Function of Bacterial Communities Emerging from Different Sources under Identical Conditions. APPL. ENVIRON. MICROBIOL. 72, 9 (2006).

26. Tseng, C.-H. et al. Microbial and viral metagenomes of a subtropical freshwater reservoir subject to climatic disturbances. ISME J 7, 2374–2386 (2013).

27. Cheaib, B. Taxon-Function Decoupling as an Adaptive Signature of Lake Microbial Metacommunities Under a Chronic Polymetallic Pollution Gradient. Frontiers in Microbiology 9, 17 (2018).

28. Findlay, S. E. G., Sinsabaugh, R. L., Sobczak, W. V. & Hoostal, M. Metabolic and structural response of hyporheic microbial communities to variations in supply of dissolved organic matter. Limnology and Oceanography 48, 1608–1617 (2003).

29. Traving, S. J. et al. Coupling Bacterioplankton Populations and Environment to Community Function in Coastal Temperate Waters. Front. Microbiol. 7, (2016).

30. Delgado-Baquerizo, M. et al. Lack of functional redundancy in the relationship between microbial diversity and ecosystem functioning. J Ecol 104, 936–946 (2016).

31. Comte, J. & del Giorgio, P. A. Links between resources, C metabolism and the major components of bacterioplankton community structure across a range of freshwater ecosystems. Environmental Microbiology 11, 1704–1716 (2009).

32. Martiny, A. C., Treseder, K. & Pusch, G. Phylogenetic conservatism of functional traits in microorganisms. ISME J 7, 830–838 (2013).

33. Jaspers, E. & Overmann, J. Ecological Significance of Microdiversity: Identical 16S rRNA Gene Sequences Can Be Found in Bacteria with Highly Divergent Genomes and Ecophysiologies. AEM 70, 9 (2004).

34. Shade, A., Chiu, C.-Y. & McMahon, K. D. Differential bacterial dynamics promote emergent community robustness to lake mixing: an epilimnion to hypolimnion transplant experiment. Environmental Microbiology 12, 455–466 (2010).

35. Wang, L.-C. et al. Increased precipitation during the Little Ice Age in northern Taiwan inferred from diatoms and geochemistry in a sediment core from a subalpine lake. J Paleolimnol 49, 619–631 (2013).

36. Tsai, J.-W. et al. Seasonal dynamics, typhoons and the regulation of lake metabolism in a subtropical humic lake. Freshwater Biology 53, 1929–1941 (2008).

37. Chen, C.-P., Tseng, C.-H., Chen, C. A. & Tang, S.-L. The dynamics of microbial partnerships in the coral Isopora palifera. ISME J 5, 728–740 (2011).

38. Jorgensen, S. L. et al. Correlating microbial community profiles with geochemical data in highly stratified sediments from the Arctic Mid-Ocean Ridge. Proceedings of the National Academy of Sciences 109, E2846–E2855 (2012).

39. Schloss, P. D. et al. Introducing mothur: Open-Source, Platform-Independent, Community-Supported Software for Describing and Comparing Microbial Communities. AEM 75, 7537–7541 (2009).

40. Edgar, R. C., Haas, B. J., Clemente, J. C., Quince, C. & Knight, R. UCHIME improves sensitivity and speed of chimera detection. Bioinformatics 27, 2194–2200 (2011).

41. Edgar, R. C. UPARSE: highly accurate OTU sequences from microbial amplicon reads. Nat Methods 10, 996–998 (2013).

42. Edgar, R. C. UNOISE2: improved error-correction for Illumina 16S and ITS amplicon sequencing. http://biorxiv.org/lookup/doi/10.1101/081257 (2016) doi:10.1101/081257.

43. Quast, C. et al. The SILVA ribosomal RNA gene database project: improved data processing and web-based tools. Nucleic Acids Research 41, D590–D596 (2012).

44. Yilmaz, P. et al. The SILVA and “All-species Living Tree Project (LTP)” taxonomic frameworks. Nucl. Acids Res. 42, D643–D648 (2014).

45. Rohwer, R. R., Hamilton, J. J., Newton, R. J. & McMahon, K. D. TaxAss: Leveraging a Custom Freshwater Database Achieves Fine-Scale Taxonomic Resolution. 3, 14 (2018).

46. Martin, M. Cutadapt removes adapter sequences from high-throughput sequencing reads. EMBnet.journal; Vol 17, No 1: Next Generation Sequencing Data AnalysisDO - 10.14806/ej.17.1.200 (2011).

47. Shen, W., Le, S., Li, Y. & Hu, F. SeqKit: A Cross-Platform and Ultrafast Toolkit for FASTA/Q File Manipulation. PLOS ONE 10 (2016).

48. Magoc, T. & Salzberg, S. L. FLASH: fast length adjustment of short reads to improve genome assemblies. Bioinformatics 27, 2957–2963 (2011).

49. Buchfink, B., Xie, C. & Huson, D. H. Fast and sensitive protein alignment using DIAMOND. Nat Methods 12, 59–60 (2015).

50. McMurdie, P. J. & Holmes, S. phyloseq: An R Package for Reproducible Interactive Analysis and Graphics of Microbiome Census Data. PLoS ONE 8, e61217 (2013).

51. Oksanen, J. et al. vegan: Community Ecology Package. (2019).

52. Wickham, H. ggplot2: Elegant Graphics for Data Analysis. (Springer-Verlag New York, 2016).

53. Parks, D. H., Tyson, G. W., Hugenholtz, P. & Beiko, R. G. STAMP: statistical analysis of taxonomic and functional profiles. Bioinformatics 30, 3123–3124 (2014).

54. J. Lin. Divergence measures based on the Shannon entropy. IEEE Transactions on Information Theory 37, 145–151 (1991).

55. Kellerman, A. M., Dittmar, T., Kothawala, D. N. & Tranvik, L. J. Chemodiversity of dissolved organic matter in lakes driven by climate and hydrology. Nat Commun 5, 3804 (2014).

56. Tanentzap, A. J. et al. Chemical and microbial diversity covary in fresh water to influence ecosystem functioning. Proc Natl Acad Sci USA 116, 24689–24695 (2019).

57. Allgaier, M. & Grossart, H.-P. Seasonal dynamics and phylogenetic diversity of free-living and particle-associated bacterial communities in four lakes in northeastern Germany. Aquat Microb Ecol 14 (2006).

58. Allgaier, M., Brückner, S., Jaspers, E. & Grossart, H.-P. Intra- and inter-lake variability of free-living and particle-associated Actinobacteria communities. Environ Microbiol 9, 2728–2741 (2007).

59. Linz, A. M. et al. Bacterial Community Composition and Dynamics Spanning Five Years in Freshwater Bog Lakes. mSphere 2, e00169–17 (2017).

60. Tandon, K. et al. Bacterial Community in Water and Air of Two Sub-Alpine Lakes in Taiwan. Microbes and environments 33, 120–126 (2018).

61. Nemergut, D. R. et al. Global patterns in the biogeography of bacterial taxa: Global bacterial biogeography. Environmental Microbiology 13, 135–144 (2011).

62. Nemergut, D. R. et al. Patterns and Processes of Microbial Community Assembly. Microbiology and Molecular Biology Reviews 77, 342–356 (2013).

63. Stegen, J. C., Lin, X., Konopka, A. E. & Fredrickson, J. K. Stochastic and deterministic assembly processes in subsurface microbial communities. The ISME Journal 6, 1653–1664 (2012).

64. Shade, A. et al. Lake microbial communities are resilient after a whole-ecosystem disturbance. ISME J 6, 2153–2167 (2012).

65. Shade, A. et al. Resistance, resilience and recovery: aquatic bacterial dynamics after water column disturbance: Bacterial community recovery after lake mixing. Environmental Microbiology 13, 2752–2767 (2011).

66. Bardgett, R. D. & Caruso, T. Soil microbial community responses to climate extremes: resistance, resilience and transitions to alternative states. Phil. Trans. R. Soc. B 375, 20190112 (2020).

67. Graham, E. B. et al. Microbes as Engines of Ecosystem Function: When Does Community Structure Enhance Predictions of Ecosystem Processes? Front. Microbiol. 7, (2016).

68. Comte, J., Fauteux, L. & Giorgio, P. A. del. Links between metabolic plasticity and functional redundancy in freshwater bacterioplankton communities. Front. Microbiol. 4, (2013).

69. Ruiz-González, C., Niño-García, J. P., Lapierre, J.-F. & del Giorgio, P. A. The quality of organic matter shapes the functional biogeography of bacterioplankton across boreal freshwater ecosystems: The functional biogeography of bacteria. Global Ecology and Biogeography 24, 1487–1498 (2015).

70. Meyer, A. F., Lipson, D. A., Martin, A. P., Schadt, C. W. & Schmidt, S. K. Molecular and Metabolic Characterization of Cold-Tolerant Alpine Soil Pseudomonas Sensu Stricto. APPL. ENVIRON. MICROBIOL. 70, 7 (2004).

71. Vick-Majors, T. J., Priscu, J. C. & A Amaral-Zettler, L. Modular community structure suggests metabolic plasticity during the transition to polar night in ice-covered Antarctic lakes. ISME J 8, 778–789 (2014).

