## Supplementary data for "Aquatic microbial community is partially functionally redundant: insights from an *in situ* reciprocal transplant experiment"

**This PDF file includes:**

Supplementary Text

Captions for Figures S1 to S8

Supplementary Fig. 1.

**Schematics of overall sample collection and data types used for analysis**. **(A)** Timeline for sample collection with the dialysis tubes incubated. **(B)** Representation of different analyses and the datatypes used to conduct the analyses.

Supplementary Fig. 2.

**16S rRNA gene-based bacterial alpha-diversity of sampling groups. (A)** Surface water richness, Shannon, Chao1, and Inverse Simpson (Inv. Simpson) diversity of different sample groups. (**B)** Bottom water richness, Shannon, Chao1, and Inverse Simpson (Inv. Simpson) diversity of different sample groups. Boxes represent the 25^th^ to 75^th^ percentiles, lines show median, and error bars represent smallest/largest values to a maximum of 1.5* Interquartile range. Statistical significance tested using Kruskal Wallis test followed by pairwise Wilcox test; * denotes *p*<0.05.

Supplementary Fig. 3.

**16S rRNA gene transcript-based alpha-diversity. (A)** Surface water richness, Shannon, Chao1, and Inverse Simpson (Inv. Simpson) diversity of different sample groups. (**B)** Bottom water richness, Shannon, Chao1, and Inverse Simpson (Inv. Simpson) diversity of different sample groups. Boxes represent the 25^th^ to 75^th^ percentiles, lines show median, and error bars represent smallest/largest values to a maximum of 1.5* Interquartile range. Statistical significance levels tested using Kruskal Wallis test followed by pairwise Wilcox test; * denotes *p*<0.05.

Supplementary Fig. 4.

16S rRNA gene based community composition and PCoA analysis for surface and bottom bacterial communities. (A–B) Bacterial community profiles at the Class level for the surface (A) and bottom (B) of both lakes were diverse and did not show significant changes when cross-swapped. Surface (C) and bottom (D) bacterial community compositions (using zero-radius Operational Taxonomic Units (zOTUs)) using the Bray-Curtis distance metric differed between lakes, even after the reciprocal transplant, with inoculum being a significant factor (*p*<0.05) for both the surface and bottom, and incubating-lake being a significant factor (*p*<0.05) driving change for bottom water only.

**Supplementary Fig. 5.**

**Ecoplate-based carbon substrate utilization.** Normalized average well color development (AWCD) as a measure of positive (>1) substrate utilization for surface samples across different sample groups.

**Supplementary Fig. 6.**

**Ecoplate-based carbon substrate utilization.** Normalized average well color development (AWCD) as a measure of positive (>1) substrate utilization for bottom samples across different sample groups.

**Supplementary Fig. 7.**

**Average well color development (AWCD) as a measure of metabolic activity of the BG community. (A)** Mean AWCD plot for surface samples from Tsuei-Feng (Ts) and Yuan-Yang Lakes (Ys) over 7 days of incubation in the laboratory. **(B)** Mean AWCD plot for bottom samples of Tsuei-Feng (Tb) and Yuan-Yang Lakes (Yb) over 7 days of incubation in the laboratory. Significant difference determined using Student’s t-test; * denotes *p<*0.05. Error bars indicate standard deviation.

**Supplementary Fig. 8.**

**Effect of average well color development (AWCD) as a measure of metabolic activity of the community with a change in the local environment. (A–B)** Yuan-Yang Lake (Y) self-swap samples (Ys(n)🡪Y; Yb(n)🡪Y) compared with cross-swap samples (Ys(n)🡪T; Yb(n)🡪T). (**C–D)** Tsuei-Feng Lake (T) self-swap samples (Ts(n)🡪T; Tb(n)🡪T) compared with cross-swap samples (Ts(n)🡪Y; Tb(n)🡪Y). Significant difference determined using Student’s t-test; * denotes *p<*0.05. Error bars indicate standard deviation. n=1,2,3 denotes sampling time after reciprocal transplant.
