## Supplementary figures and images for "Aquatic microbial community is partially functionally redundant: insights from an *in situ* reciprocal transplant experiment"

### Supplementary Figure S1

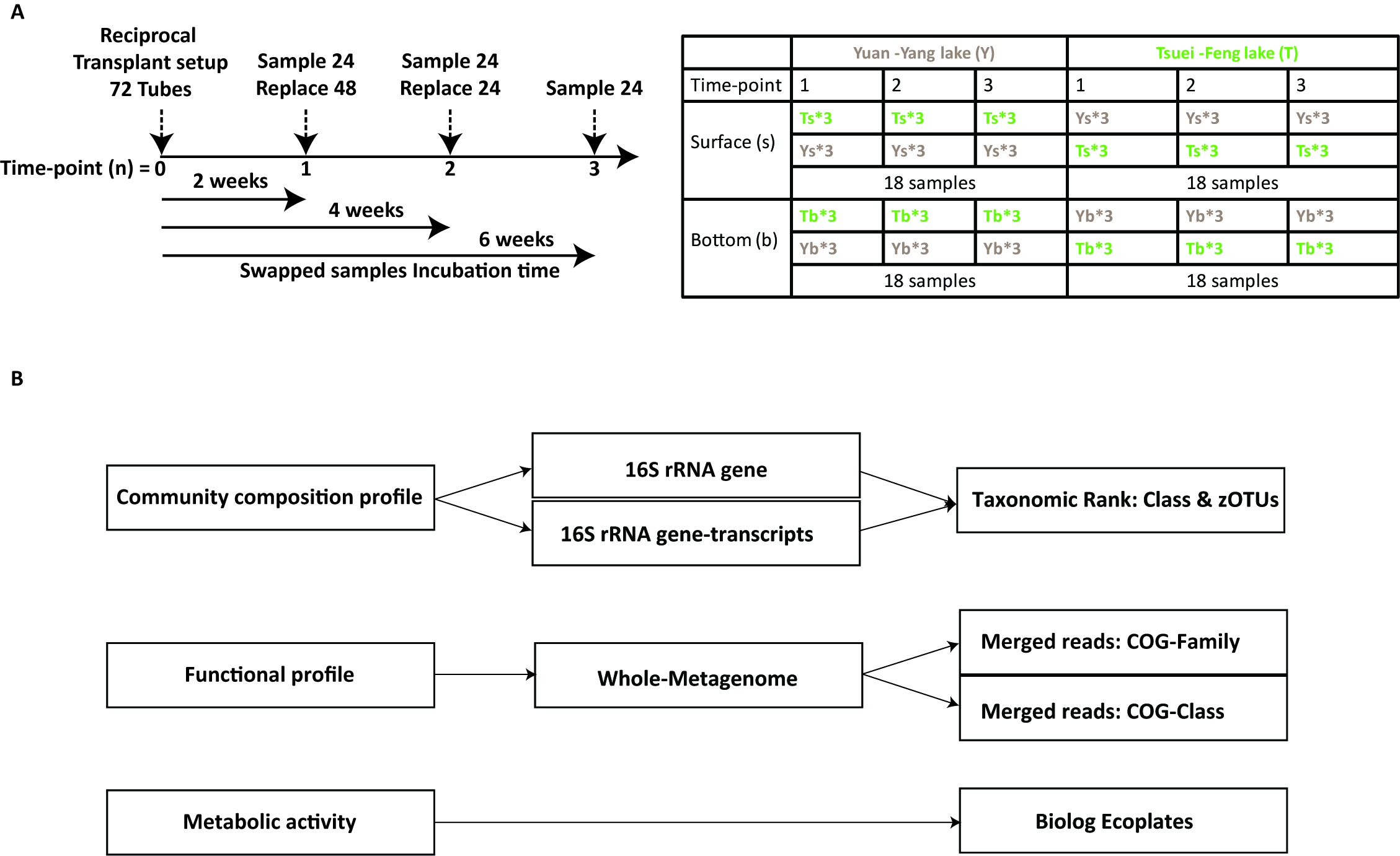

### Supplementary Figure S2

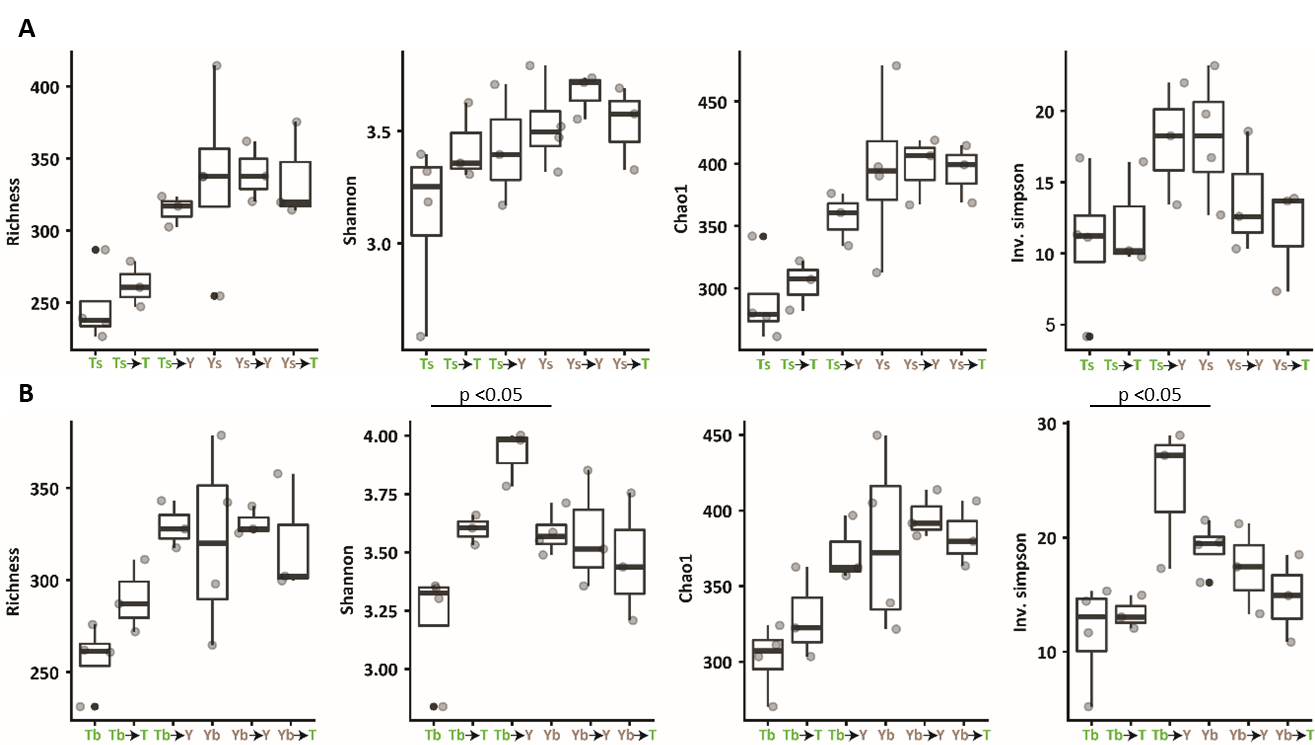

### Supplementary Figure S3

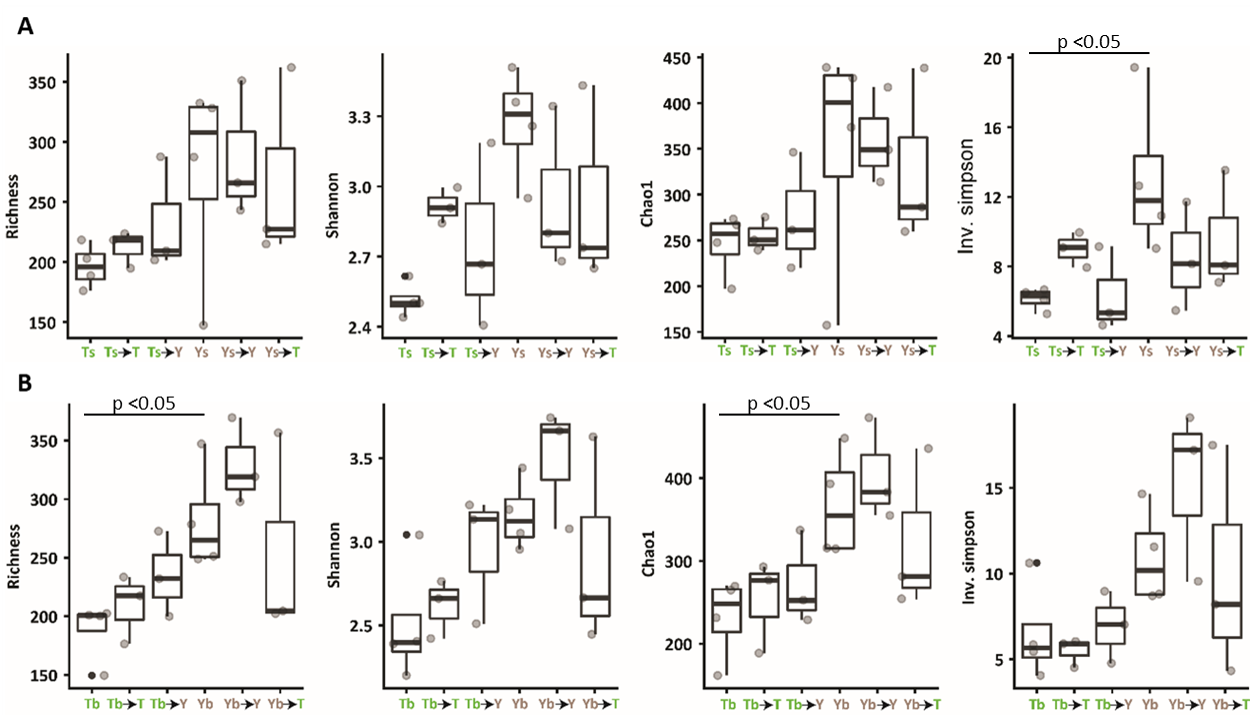

### Supplementary Figure S4

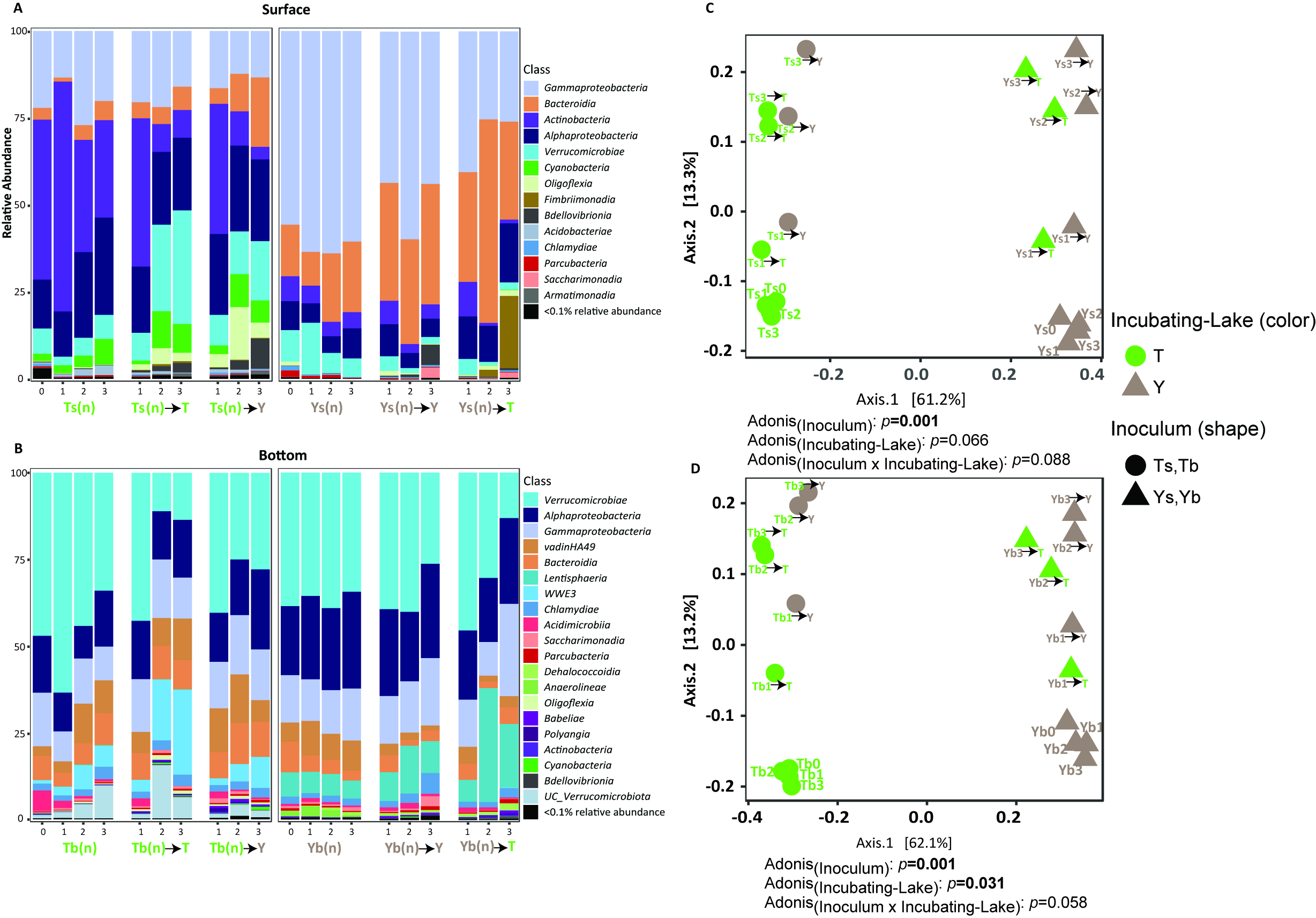

### Supplementary Figure S5

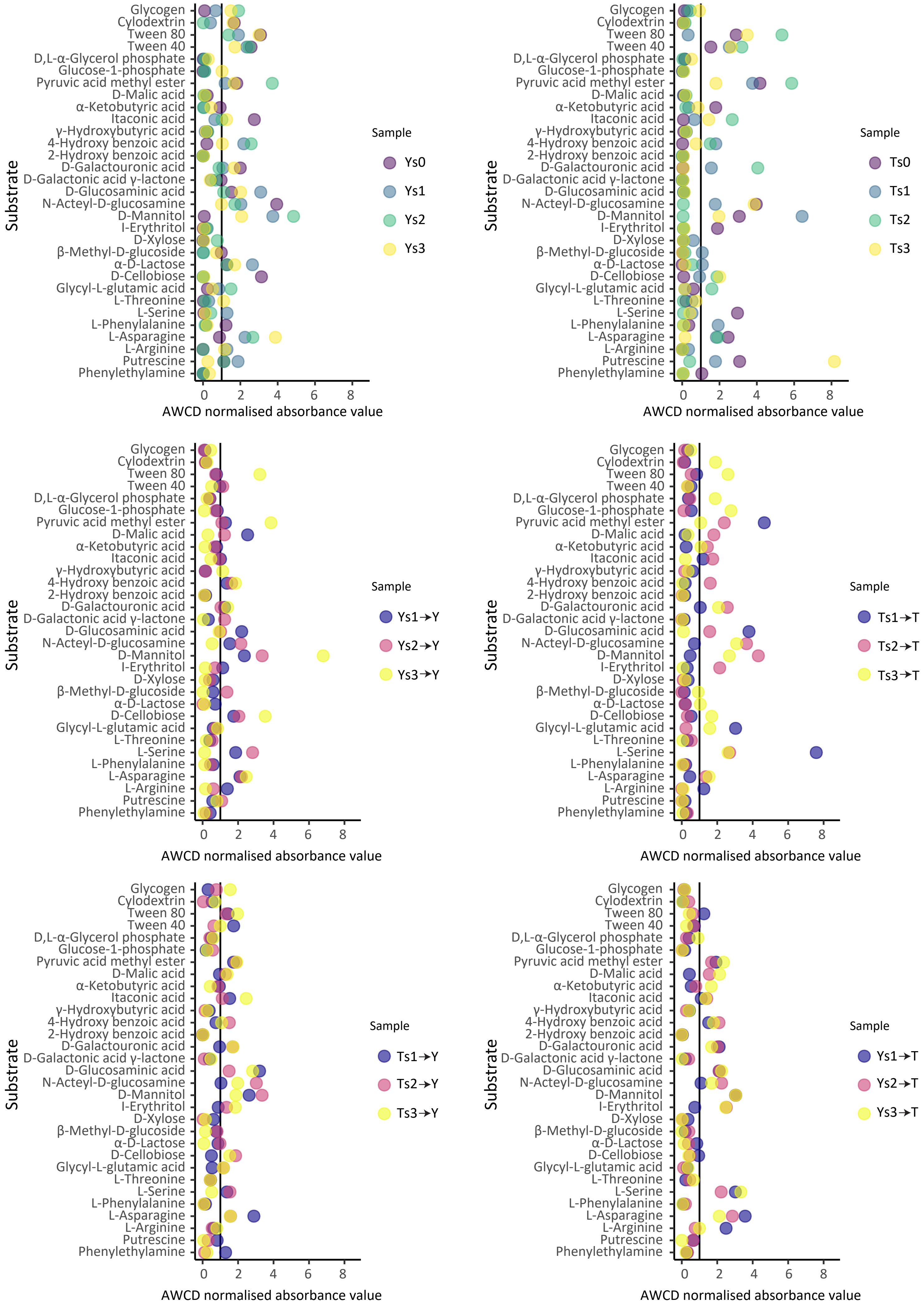

### Supplementary Figure S6

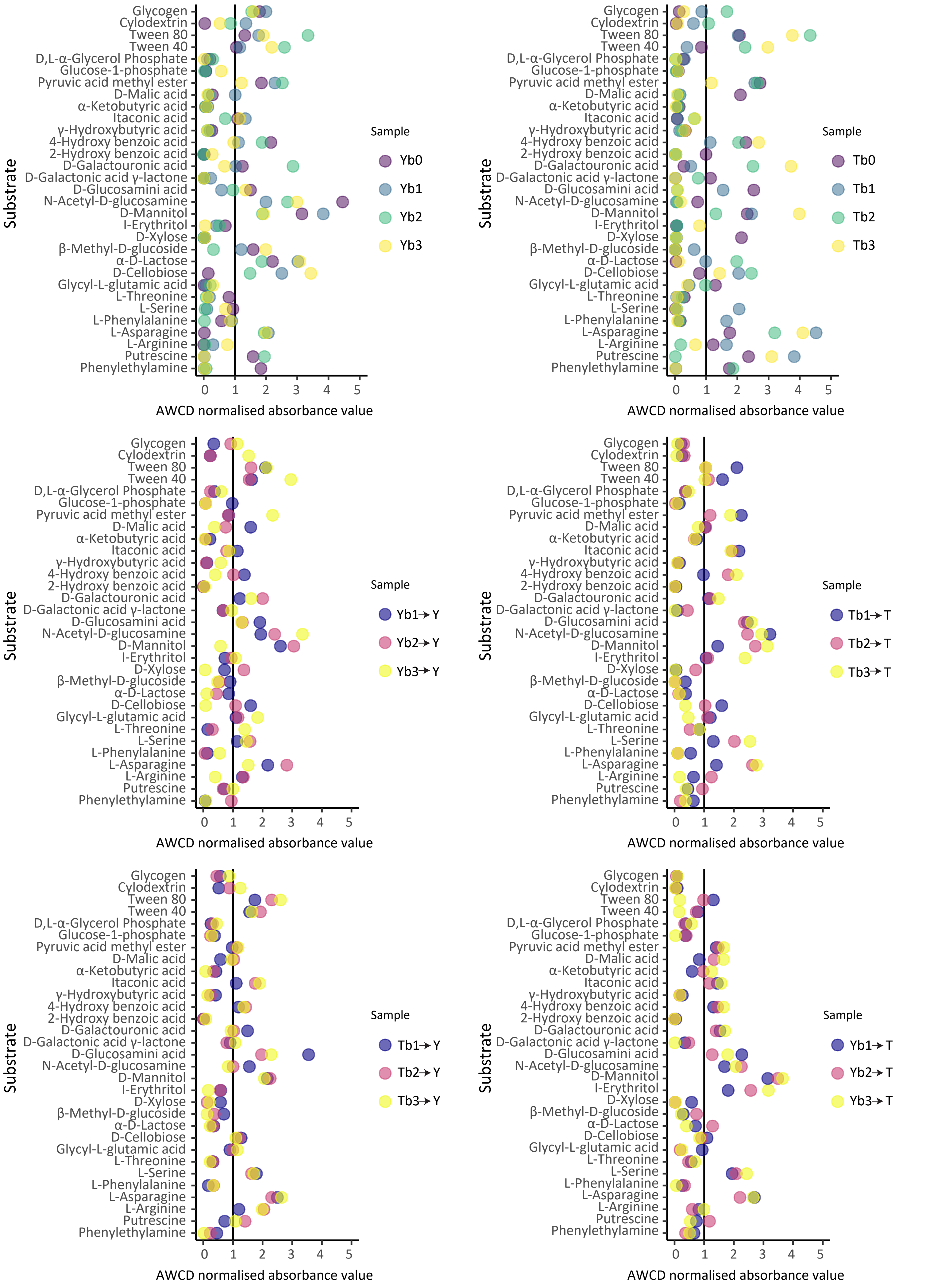

### Supplementary Figure S7

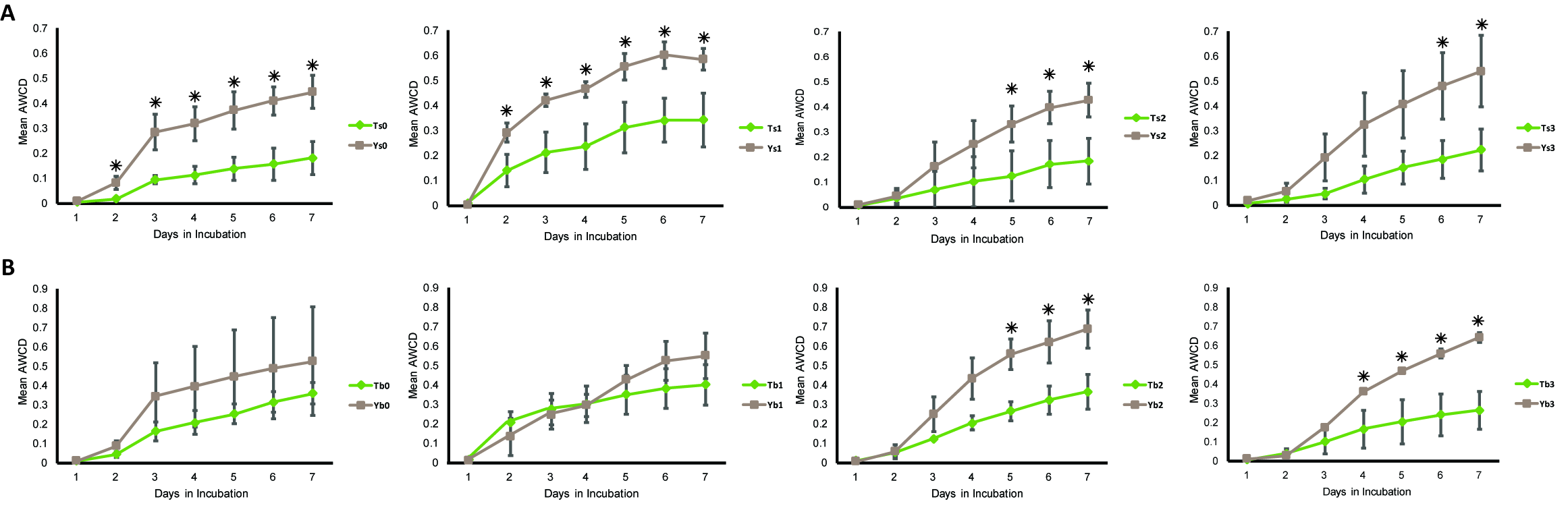

### Supplementary Figure S8

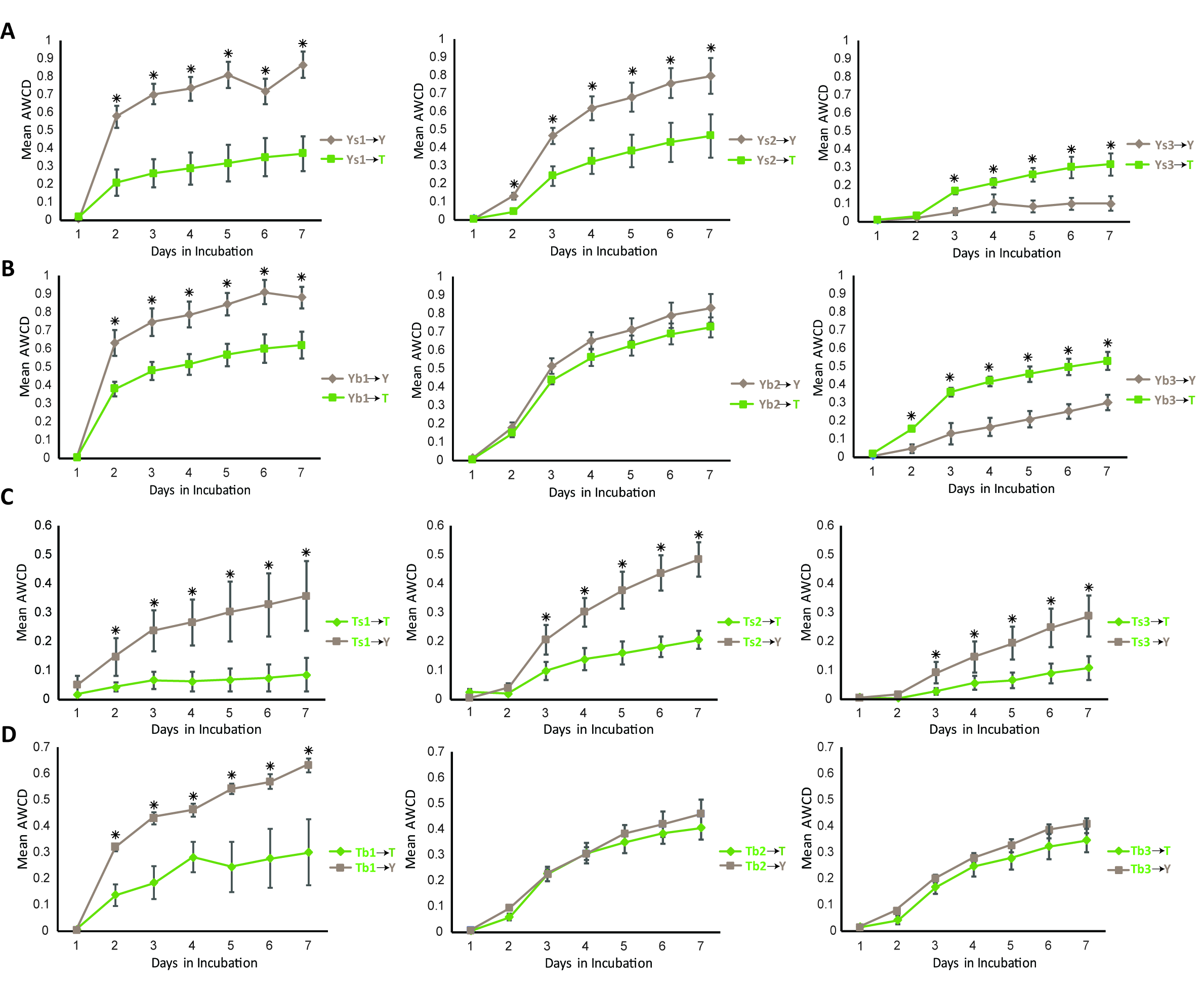
